# Histone Variant H2A.J Links Epigenetic Reprogramming to Mitochondrial-dependent Kidney Regeneration under Radiation Stress

**DOI:** 10.64898/2026.06.18.733158

**Authors:** Mutaz Abd Al-razaq, Julia von der Lippe, Nicolas Freche, Daniela Jung, Marlene Jordan, Henrik Auerbach, Markus Hecht, Christian Rübe, Daniela Kramer, Carl Mann, Claudia E. Rübe

## Abstract

The histone variant H2A.J is implicated in radiation-induced senescence by promoting the transcription of inflammatory genes. While H2A.J expression increases in renal tubular epithelial cells (TECs) following ionizing radiation (IR), its functional role remains poorly understood. To investigate this, constitutive H2A.J knock-out (KO) mice and wild-type (WT) controls were subjected to CT-guided IR (fractionated whole-body or localized kidney IR). Kidneys were analyzed at acute, intermediate, and chronic stages using immunofluorescence, histochemistry, automated image analysis, and electron microscopy. In WT TECs, IR induced rapid chromatin incorporation and C-terminal serine phosphorylation of H2A.J. Conversely, KO kidneys exhibited significantly more severe histopathological damage, including tubular dilation, flattened epithelium, associated with increased apoptosis, and premature senescence, characterized by persistent DNA damage with lamin B1 loss. Notably, KO TECs displayed disrupted mitochondrial networks and reduced brush borders even at baseline, which were further exacerbated by IR. Unlike WT controls, KO kidneys developed progressive tubular atrophy and incipient fibrosis, indicating a failure in regenerative capacity. Our findings suggest that H2A.J loss impairs tubular regeneration due to defective mitochondrial activation, resulting in insufficient energy supply for coordinated repair. Collectively, these results identify H2A.J as a critical stress-adaptive histone variant essential for the epigenetic regulation of tissue repair following radiation-induced damage.

**One Sentence Summary:** In irradiated kidney, the loss of histone variant H2A.J impairs the chromatin-mediated adaptation of mitochondrial function in tubular epithelial cells, thereby exacerbating cellular stress - characterized by increased induction of apoptosis and senescence - and ultimately leading to tubular atrophy.

**Translational Relevance:** Acute and chronic kidney injury are frequent complications of genotoxic cancer therapies. Chemo- and radiotherapy induce DNA lesions that trigger cell death and senescence, often leading to irreversible renal damage. However, renal regeneration can occur through the dedifferentiation, proliferation, and redifferentiation of surviving tubular epithelial cells (TECs). This repair process is governed by epigenetic mechanisms that regulate the DNA damage response (DDR) and adapt gene expression programs. Following ionizing radiation (IR), epigenetic remodeling involves the incorporation of histone variants that modulate chromatin accessibility for stress-responsive transcription factors. We identify the histone variant H2A.J as a constitutive component of renal TECs, significantly upregulated after exposure to ionizing radiation (IR). Using H2A.J knock-out (KO) mice, we demonstrate that its absence disrupts acute damage responses and prevents coordinated repair, severely impairing regeneration. Mechanistically, H2A.J deficiency compromises mitochondrial function under postirradiation metabolic stress, driving the transition from acute injury to chronic kidney disease via persistent inflammation and maladaptive tubulointerstitial repair. Targeting these epigenetic drivers offers a promising strategy for regenerating damaged kidney tissue in oncology.

## INTRODUCTION

Kidney damage caused by ionizing radiation (IR) is a serious side effect of radiotherapy for tumors in the upper abdominal cavity. Following radiation-induced damage both acute injury and chronic kidney disease are pathophysiologically linked syndromes, so-called radiogenic nephropathy, which can ultimately lead to complications such as end-stage renal failure (1). The occurrence of clinically manifest renal dysfunction following IR exposure is highly dependent on cumulative dose, dose per fraction and irradiated kidney volume (2,3).

Genotoxic stress resulting from IR exposure leads to lesions in nuclear and mitochondrial DNA and, consequently, to the activation of the DNA damage response (DDR) - such as DNA repair, reversible cell cycle arrest, or premature senescence - as well as to various forms of cell death (apoptosis, necroptosis, etc.) (4). Following acute kidney injury, the most severely affected renal structures are the cortical compartments, with tubular epithelial cells (TECs) being especially vulnerable due to their high energetic and metabolic demands (5). Along the nephron, TECs in the proximal tubules are responsible for reabsorption of glomerular filtrates, which requires continuous active ion transport against steep concentration gradients. Due to their constantly high ATP consumption, TECs require large numbers of mitochondria to maintain basal homeostasis, and in particular, to recover successfully after acute damage (6). Tubular epithelium regenerates through a highly coordinated sequence of events, whereby surviving TECs dedifferentiate, re-enter cell cycle, and proliferate to replace lost or damaged epithelial cells (7–9). Newly proliferated cells migrate along the basement membrane to cover deepithelialized areas and restore the epithelial barrier (10). Once coverage is complete, these tubular cells redifferentiate, thereby regaining their apical-basolateral polarity, brush border structures and functional capacity (11). Since TECs are highly energy-dependent, maintaining mitochondrial function is essential to adapt energy metabolism as needed for effective regeneration after acute kidney damage, thus enabling the restoration of normal tubular structure and function (6). However, if functional adaption of mitochondria to cellular stress is impaired, nephrotoxic processes can become maladaptive, resulting in chronic inflammation and ultimately renal fibrosis (12,13).

In cell nuclei, DNA is wrapped around histone octamers forming nucleosomes and thereby contributing to the organization and compaction of genetic material. Histone modifications and histone variants reorganize the nuclear landscape in senescent cells (14,15), but are also essential for acute stress responses; by rapidly remodeling chromatin structures, they enable the transcriptional flexibility required to activate survival and repair pathways, thereby determining cell fate following damage (16). This epigenetic reorganization is essential for TECs to rapidly switch between injury, repair, and differentiation states, precisely controlling which genes are turned on or off during the regeneration process (17). Epigenetic regulation at the level of histones may therefore fundamental for controlling how TECs respond to and recover from acute injury.

The histone variant H2A.J has been identified in connection with replicative and premature senescence resulting from genotoxic stress – e.g. the induction of DNA double-strand breaks (DSBs) - whereby it modulates the transcription of proinflammatory mediators as part of the senescence-associated secretory phenotype (SASP) (18). H2A.J differs only slightly in its amino acid sequence from canonical H2A; however, it possesses a phosphorylation site at the nucleosome-H1 interface, and can therefore weaken H1-nucleosome interaction, decompact chromatin, and increase DNA accessibility (19,20). This strategic location allows a single modification to reshape chromatin from the nucleosome level up to large-scale chromatin domains. The enrichment of H2A.J in glandular and epithelial tissues fits its role as an "epigenetic stress-sensor" that tunes the inflammatory response of barrier tissues to environmental damage (18,21,22). Previous in-vitro experiments in human fibroblasts showed that differential incorporation of H2A.J after IR exposure significantly affects global chromatin organization, recruitment of transcription factors, and consequently secretion of inflammatory cytokines (23,24). In a mouse model involving skin IR, H2A.J is upregulated and modulates - acting as a driver of SASP gene expression - acute and chronic inflammation within the context of complex tissue responses (25).

Since H2A.J is increasingly expressed in renal TECs within hours following IR, these preclinical studies aimed to investigate the pathophysiological significance of H2A.J in the context of radiation-induced nephropathy using a constitutive H2A.J knockout mouse model.

## RESULTS

### H2A.J expression and DSB repair in aging kidneys (Fig. 1)

Histologically, the renal parenchyma comprises the cortex, containing glomeruli and convoluted tubules, and the medulla, organized into pyramids of straight tubules and collecting ducts. Since H2A.J is associated with replicative senescence - and thus with the potential aging of organs - we examined kidney tissue from young (2–3 months), middle-aged (12 months), and aged (18 months) WT mice. With age, the renal cortex exhibits structural decline, including a reduction in functional glomeruli, compensatory glomerular hypertrophy (increasing from ∼60 µm to ∼80 µm), tubular atrophy, and luminal dilation (Fig. 1A) (26). Notably, H2A.J expression in TECs increased age-dependently, confirming our previous findings (Fig. 1A) (18).

**Fig. 1:**
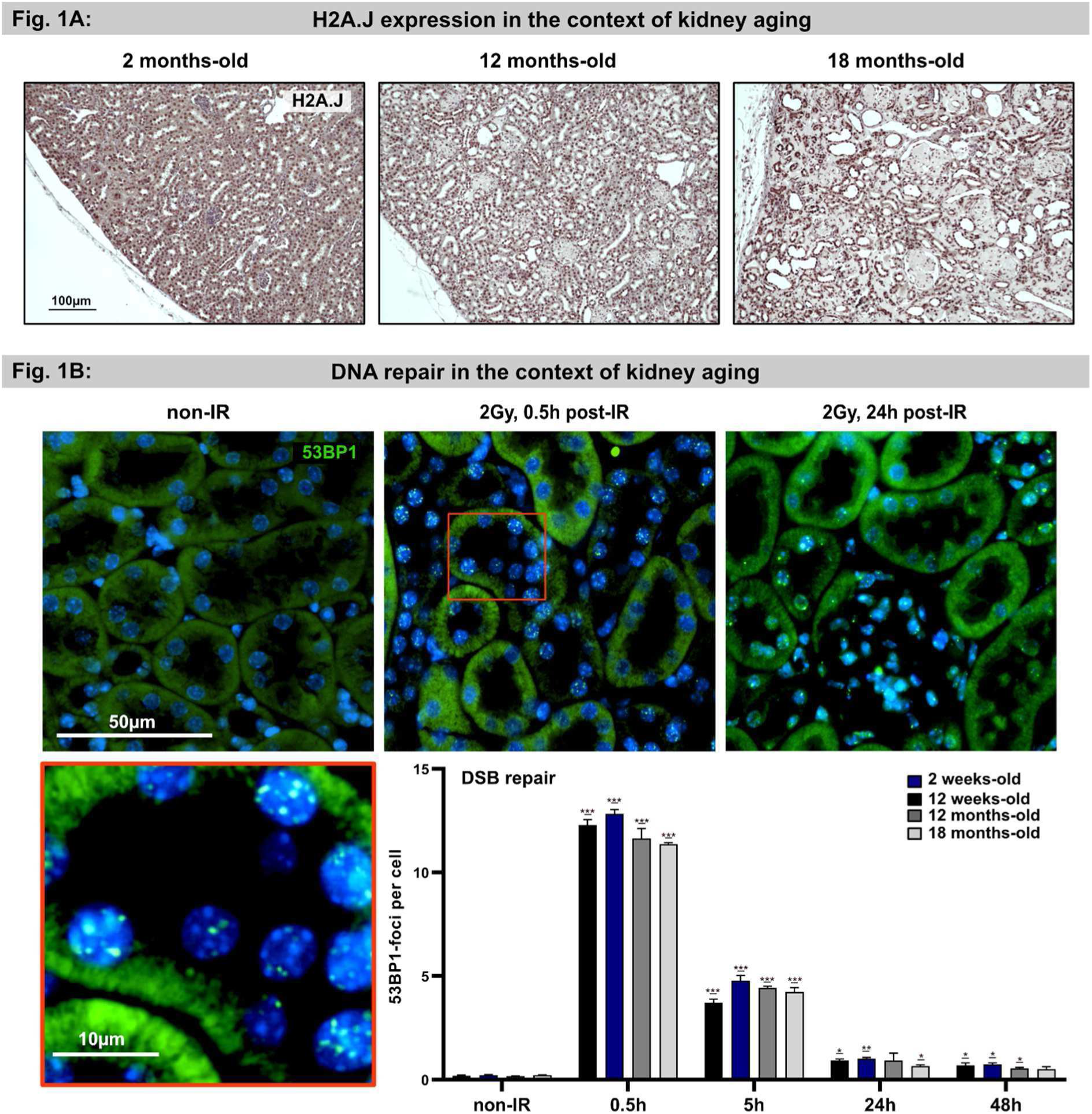
H2A.J in the context of physiological aging of WT kidneys. A: Representative micrographs of H2A.J-stained kidney from wild-type (WT) mice across different age groups. B: Micrographs of 53BP1-stained kidney at 0.5 h and 24 h post-IR compared to non-irradiated controls. Quantification of 53BP1-foci per nucleus indicates DNA double-strand break (DSB) repair kinetics as a function of age. Data are presented as mean ± SD (n ≥ 3). Significant deviations from non-irradiated controls are indicated by asterisks. P-values were defined as: *p < 0.05, **p < 0.01, and ***p < 0.001.

As histone H2A variants are critical for DSB signaling and repair (27), we assessed age-dependent DSB repair efficiency using 53BP1-foci kinetics after 2 Gy. Maximum foci induction occurred at 0.5 h (12.3 ± 0.2 foci/nucleus), followed by a marked reduction at 5 h (3.7 ± 0.2 foci/nucleus) and near-complete repair within 24 h post-IR (0.9 ± 0.1 foci/nucleus) (Fig. 1B). No significant differences were observed across age groups, suggesting that DSB repair efficiency in the kidney remains stable throughout aging; furthermore no significant increase in basal 53BP1-foci was detected at 18 months.

### H2A.J expression in WT versus KO kidney following IR exposure (Fig. 2)

To investigate radiation-induced genotoxic stress, WT and KO kidneys were analyzed following fractionated (5x 2 Gy) or high-dose (10 Gy) IR (Suppl. 1). In WT kidneys, H2A.J expression was predominantly localized to cortical tubular nuclei, with minimal staining in glomeruli or collecting ducts. Automated quantification revealed a dose-dependent increase in H2A.J+ cells in WT parenchyma, rising from 23.1 ± 9.7% (non-IR) to 36.3 ± 16.7% (5x 2 Gy, 72 h post-IR) and 62.8 ± 5% (10 Gy, 24 h post-IR), with expression persisting for at least one week. Using a phospho-specific antibody, we detected H2A.J phosphorylation within 24 hours post-IR (Suppl. 2). Given its position at the nucleosome-H1 linker junction, H2A.J phosphorylation likely modulates chromatin architecture and nucleosome dynamics, potentially facilitating access for transcription factors and remodeling complexes during the DDR. As expected, H2A.J was undetectable in KO kidneys (Suppl. 3).

**Fig. 2:**
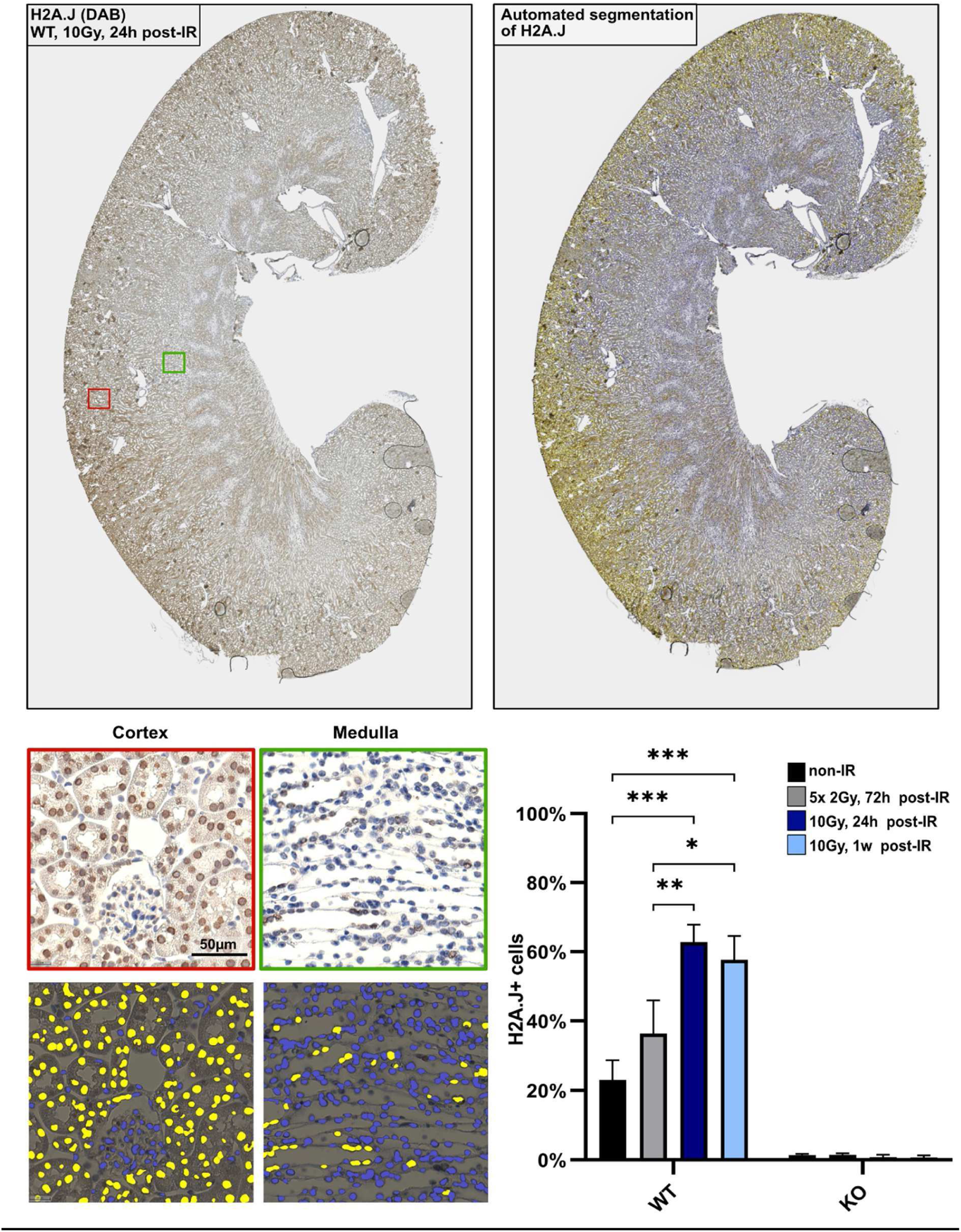
Quantification of H2A.J+ cells in WT and KO kidneys following IR exposure. Automated segmentation of H2A.J+ cells in wild-type (WT) and knock-out (KO) kidneys under baseline conditions and following IR. Data are presented as mean ± SD (n ≥ 3). Group differences were evaluated via oneway or two-way ANOVA. P-values were defined as: *p < 0.05, **p < 0.01, and ***p < 0.001.

### Early histopathological alterations and premature senescence (Fig. 3)

Within 24 hours following IR exposure, histopathological changes are detectable in WT but clearly more pronounced in KO kidneys, indicating the onset of severe cellular stress, particularly in cortical compartments. Genotoxic stress triggers excessive reactive oxygen species (ROS) and mitochondrial DNA (mtDNA) damage, causing metabolic failure and ATP depletion that impairs Na+/K+-ATPase function, leading to cellular swelling and subsequent interstitial edema. In irradiated KO kidneys, proximal tubules already show significant epithelial flattening, whereas glomerular structure remains largely intact (Fig. 3A), suggesting that TECs are losing polarity and expanding to close epithelial gaps. Since persistent DNA damage signals often trigger premature senescence, 53BP1-foci and nuclear lamin B1 levels were investigated. At 24 h following IR exposure, WT kidneys show healthy tubular epithelium with strong nuclear H2A.J and lamin B1 signals, and minimal persistent 53BP1-foci (Fig. 3B). In contrast, KO kidneys exhibit flattened epithelium, no H2A.J and reduced Lamin B1 signals (Suppl. 4), and numerous persistent 53BP1-foci were observed (Fig. 3B). Collectively, these results suggest that absence of H2A.J may lead to a disrupted DDR with increased senescence induction in KO TECs.

**Fig. 3:**
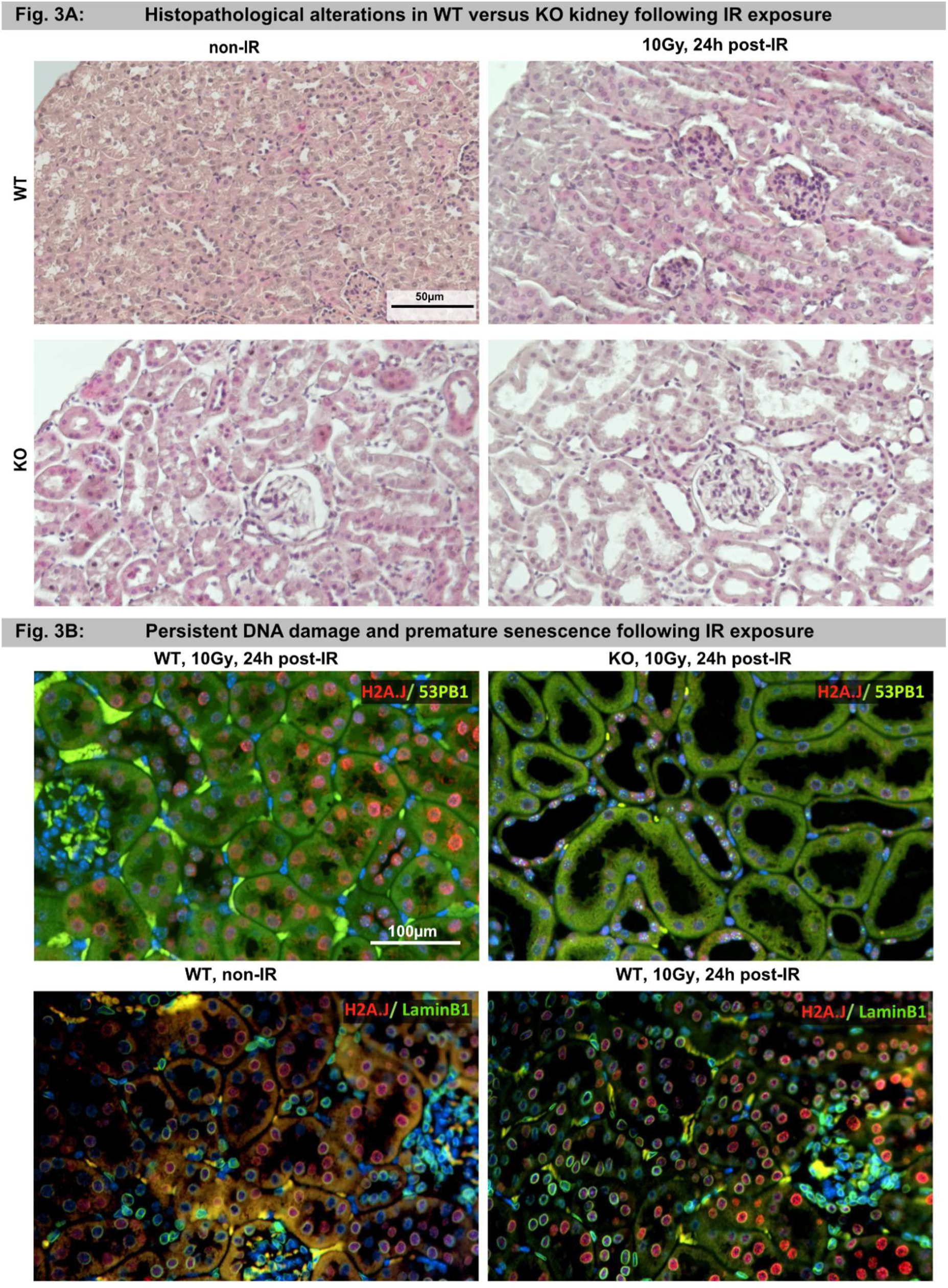
Histopathological alterations in WT and KO kidneys following IR exposure. A: Micrographs of H&E-stained WT and KO kidneys before (non-IR) and after IR exposure (10Gy, 24h post-IR). B: Micrographs of IF-stained (H2A.J, 53BP1, Lamin B1) of WT and KO before and after IR exposure to assess premature senescence.

### DNA damage response in WT versus KO kidney following IR exposure (Fig. 4)

Following genotoxic stress, the DDR acts as a molecular switchboard that determines cell fate based on the severity of damage. Sustained DDR signaling can drive cells into premature senescence, often characterized by persistent DNA damage foci. To compare the amount of senescence induction in WT versus KO kidneys, cells with 53BP1-foci were quantified at 24 h following IR exposure with 10 Gy.

**Fig. 4:**
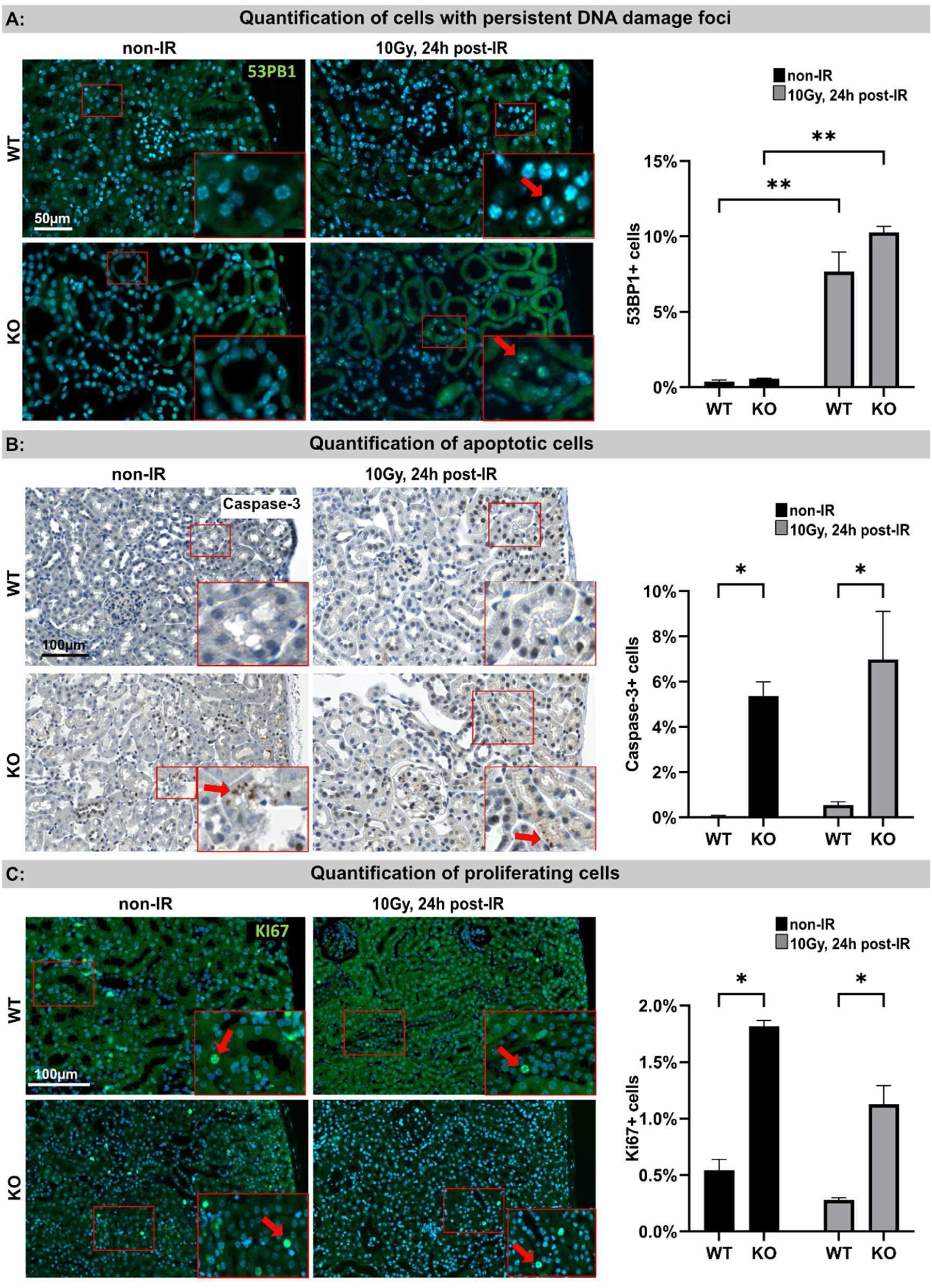
DNA damage response in WT and KO kidneys following IR exposure. A: Micrographs of 53BP1-stained WT and KO cortex at 24 h post-IR compared to non-irradiated controls. Quantification of cells with persistent foci. B: Micrographs of Caspase-3-stained WT and KO cortex at 24 h postIR compared to non-irradiated controls. Quantification of apoptotic cells. C: Micrographs of Ki67-stained WT and KO cortex at 24 h post-IR compared to non-irradiated controls. Quantification of proliferating cells. Data are presented as mean ± SD (n ≥ 3). P-values were defined as: *p < 0.05 and **p < 0.01.

While baseline levels in non-irradiated renal parenchyma were low, 53BP1+ cells increased significantly in WT (7.7 ± 1.3%) and even more in KO kidneys (10.3 ± 0.4%) at 24 h post-IR (Fig. 4A). When radiation-induced DNA damage is too severe, IR triggers programmed cell death via the proteolytic activation of executioner caspases, such as caspase-3, to rapidly and effectively eliminate injured cells without eliciting an inflammatory response. In renal parenchyma, apoptosis rates (quantified by caspase-3+ cells) were markedly higher in KO kidneys, reaching 5.4 ± 0.6% at baseline and 6.9 ± 2.1% post-IR, compared to <0.5 ± 0.1 % in WT (Fig. 4B). In tubular epithelium, this programmed cell death represents a direct response to excessive genotoxic stress and mitochondrial damage, ensuring that severely compromised cells are eliminated rather than persisting in a dysfunctional state.

In healthy kidneys, proliferation is minimal as most renal cells - particularly within proximal tubules and glomeruli - remain in a quiescent state due to very low cellular turnover. However, following injury, the loss of tubular cells through senescence or apoptosis forces surviving cells to proliferate to close epithelial gaps and prevent organ failure. Notably, non-irradiated KO kidneys displayed significantly higher baseline proliferation (Ki67+ cells: 1.8 ± 0.1%) compared to WT (0.5 ± 0.1%), a trend that persisted post-irradiation despite a slight overall decline in counts (Fig. 4C). During acute radiation injury, surviving TECs must dedifferentiate and proliferate to restore epithelial integrity (28). In KO kidneys, the absence of H2A.J triggers premature senescence and increased apoptosis, necessitating this elevated proliferation to maintain tissue homeostasis.

Notably, Prussian blue staining (for the detection of hemosiderin deposits) revealed impaired tubular function with disrupted iron reabsorption—not only in irradiated kidneys but also in KO kidneys prior to IR, suggesting a fundamental impairment of tubular performance (Suppl. 5).

### Mitochondrial stress response in WT versus KO kidney following IR exposure (Fig. 5)

IR triggers reactive oxygen species (ROS) production, leading to mitochondrial damage, impaired oxidative phosphorylation, and energy failure. To assess mitochondrial integrity, we stained for Translocase of Outer Mitochondrial Membrane 20 (TOMM20), a marker generally used to visualize mitochondrial networks within tissue cells. In non-irradiated WT kidneys, TOMM20 staining was intense and homogenous (73.6 ± 3.3%), particularly in cortical TECs. However, 24 hours post-IR (10Gy), we observed a significant reduction in mitochondrial mass (47.3 ± 11.4%) likely due to increased oxidative stress (Fig. 5). In contrast, non-irradiated KO kidneys displayed lower TOMM20 levels with a distinct, patchy distribution at the luminal pole - a pattern exacerbated by IR (Fig. 5B). Such irregular or fragmented TOMM20 staining pattern typically indicate disturbed mitochondrial network integrity and impaired mitochondrial function, including altered oxidative phosphorylation (29).

**Fig. 5:**
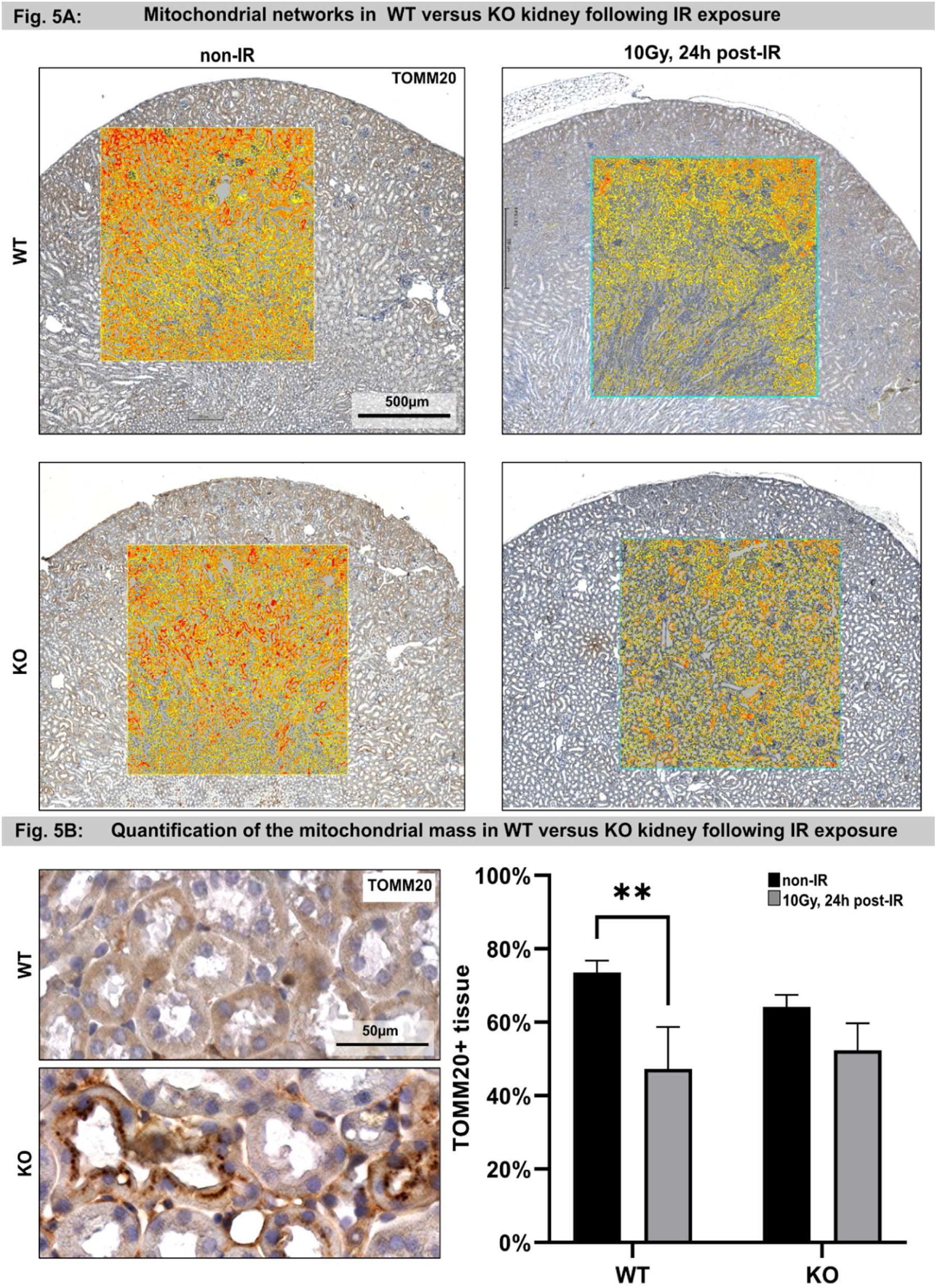
Mitochondrial networks in WT and KO kidneys following IR exposure. A: Representative micrographs of TOMM20-stained WT and KO kidney at 24 h post-IR compared to non-irradiated controls. Insets show automated image segmentation used to visualize staining intensity. B: Automated image analysis-based quantification of TOMM20+ tissue area. Data are presented as mean ± SD (n ≥ 3). P-values were defined as: **p < 0.01.

### Ultrastructural alterations in TECs of WT versus KO kidney following IR exposure (Fig. 6)

Radiation-induced mitochondrial damage in renal TECs can impair cellular energy metabolism and cytoskeletal integrity, which in turn can contribute to the loss of their specialized phenotype with reduction of apical brush border structures (Suppl. 6). To achieve a direct visualization of ultrastructural damage at nanoscale resolution, we conducted electron microscopic investigations of the renal cortex. In WT kidneys, Transmission Electron Microscopy (TEM) images of proximal tubules showed normal nuclei, abundant mitochondria, and dense brush borders. In contrast, non-irradiated KO TECs already exhibited patchy heterochromatin, smaller mitochondria, and shortened microvilli (Fig. 6A). Three months post-IR, both mouse lines showed increased chromatin condensation, though damage was markedly more severe in KO kidneys (Fig. 6A). While irradiated WT kidneys displayed mitochondrial shrinkage and microvilli reduction, KO TECs progressed to near-complete destruction, characterized by ruptured cytoplasm and absent brush borders. Scanning Electron Microscopy (SEM) analysis corroborated these findings, demonstrating intact tubular structures in non-irradiated WT kidneys. In contrast, irradiated WT kidneys exhibited luminal obstruction by cellular debris (Fig. 6B). Notably, non-irradiated KO kidneys already presented with collapsed lumens, while irradiated KO kidneys showed extensively destroyed tubules characterized by massive tissue remodeling. Ultimately, these ultrastructural changes demonstrate that the absence of H2A.J exacerbates radiation-induced damage, leading to profound loss of tubular integrity and functionality.

**Fig. 6:**
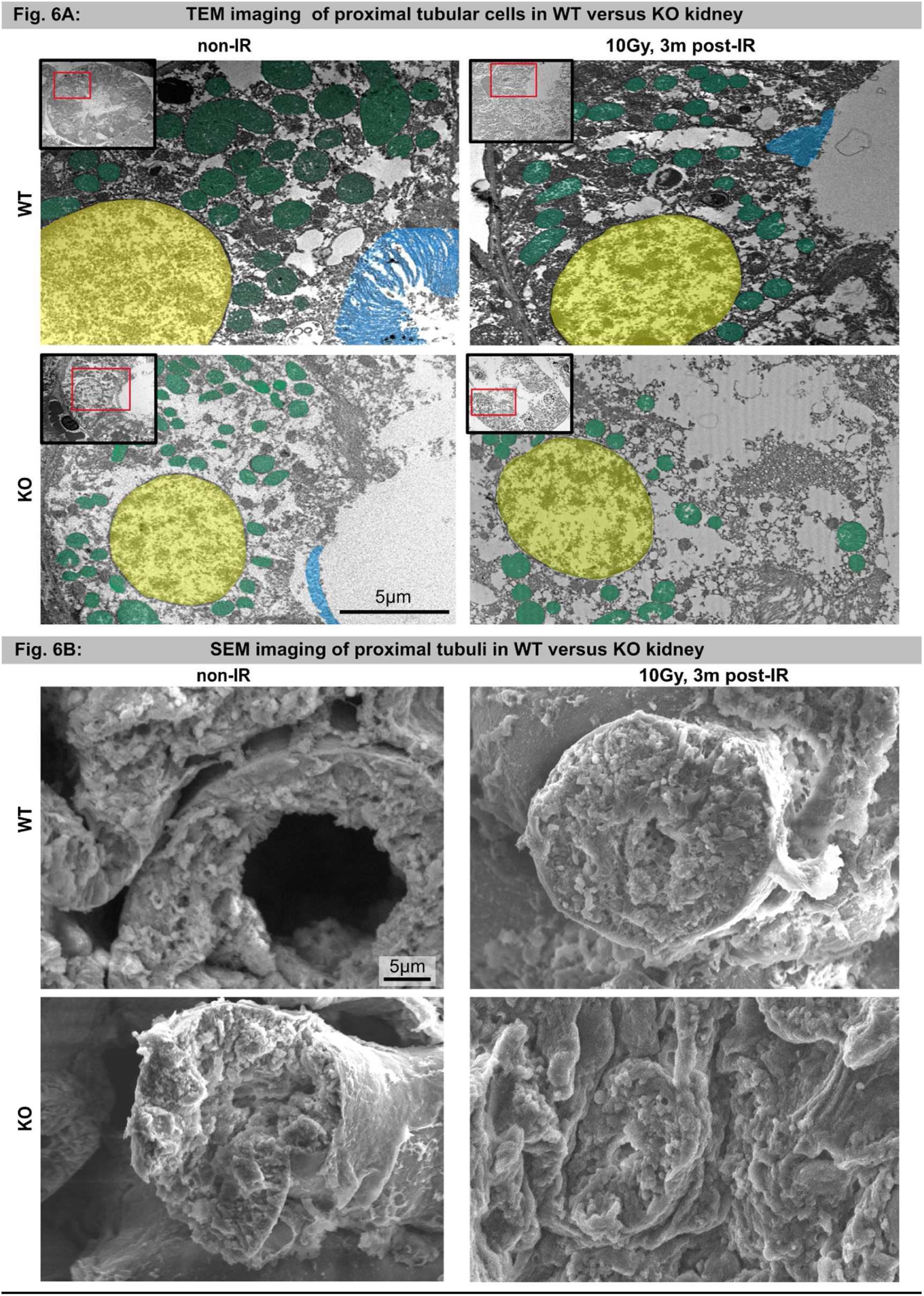
Electron microscopy of proximal tubules in WT and KO kidneys following IR exposure. A: TEM imaging of TECs in WT and KO kidneys at 3 months post-IR compared to non-irradiated controls. Nuclei, mitochondria, and the brush border are pseudo-colored in yellow, green, and blue, respectively. B: SEM imaging of proximal tubules in WT and KO kidneys at 3 months post-IR compared to non-irradiated controls.

### Tubulus atrophy and incipient fibrosis in KO kidney following IR exposure (Fig. 7)

To investigate the long-term effects of IR, Masson’s trichrome staining was used to assess structural remodeling and renal fibrosis. While irradiated WT kidneys showed only minor histological changes, irradiated KO kidneys exhibited pronounced tubular atrophy - characterized by luminal narrowing - and increased peritubular collagen accumulation (Fig. 7). The specific distribution of incipient fibrosis in the tubulointerstitial space, rather than periglomerular regions, identifies the proximal tubules as the primary site of chronic injury. These findings suggest that irreversible damage to TECs facilitates the transition from acute to chronic kidney injury. Furthermore, aged and irradiated KO mice - but not WT controls - developed progressive tubular atrophy and renal cysts (Suppl. 7). This cystic transformation likely results from a loss of epithelial differentiation with impaired tubular transport functions, leading to fluid accumulation, luminal dilation, and the progressive development of renal cysts (Suppl. 8) (30).

**Fig. 7:**
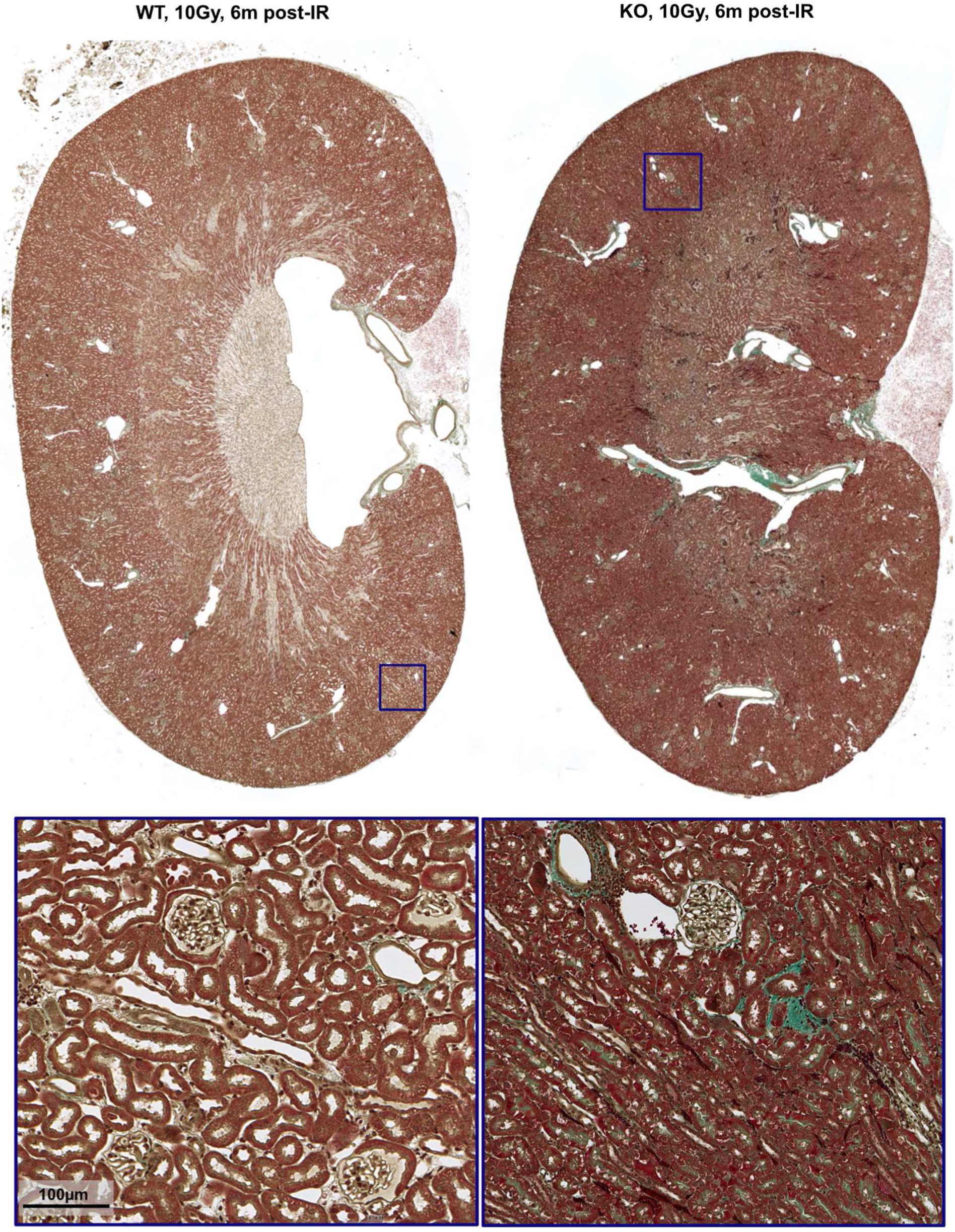
Tubular atrophy and incipient fibrosis in KO kidneys following IR exposure. Micrographs of Masson-Goldner-stained WT and KO kidneys at 6 months post-IR. Insets provide higher magnification highlighting pronounced tubular atrophy in KO kidney.

## DISCUSSION

This study demonstrates that radiation-induced genotoxic stress triggers rapid H2A.J incorporation and phosphorylation within the TECs of WT kidneys. While radiation-induced tissue alterations are effectively mitigated by adaptive mechanisms in WT tissues, H2A.J-deficient kidneys exhibit a markedly divergent phenotype driven by epigenetic dysregulation. In the absence of H2A.J, the synergistic impact of increased cell death and premature senescence drives a pro-inflammatory milieu. Subsequent maladaptive compensatory proliferation, coupled with a failure to redifferentiate into functional tubular epithelia, promotes progressive tissue remodeling and exacerbates tubular atrophy. We hypothesize that the epigenetic regulation of mitochondrial function in H2A.J-deficient kidneys is fundamentally impaired - a deficit that is markedly exacerbated following IR exposure. This suggests that H2A.J-mediated epigenetic regulation of mitochondrial homeostasis is indispensable for meeting the high metabolic demands of cellular recovery after genotoxic stress. Collectively, H2A.J appears to provide the requisite epigenetic plasticity to coordinate adaptive stress responses in TECs by ensuring sufficient bioenergetic capacity, thereby maintaining tubular integrity.

### Histone variants for reprogramming chromatin states in response to genotoxic stress

Cells respond to genotoxic stress by replacing canonical histones with specific variants to remodel chromatin accessibility and modulate transcriptional programs (31). This epigenetic reprogramming is essential for transitioning from the initial DDR to a coordinated phase of tissue regeneration. In proximal TECs, H2A.J serves as a pivotal epigenetic regulator of these differentiation transitions. Specifically, the phosphorylation of unique serine residues can act as a signal-responsive regulatory switch, capable of rapidly modulating higher-order chromatin architecture (32). Conversely, dysfunctional H2A.J signaling appears to compromise the renal regenerative response and sustain chronic inflammation, thereby predisposing the tissue to chronic kidney disease progression. Previous studies have demonstrated that H2A.J promotes the secretion of SASP cytokines via the activation of NF-κB and STAT signaling, with its dysregulation profoundly altering the inflammatory milieu (18,33). Furthermore, H2A.J has been shown to influence cell cycle progression and epithelial-mesenchymal transition (EMT) in response to IR (23,34). By modulating cellular plasticity - specifically the dedifferentiation and transdifferentiation of TECs - H2A.J appears to be indispensable for an orchestrated regenerative response.

### H2A.J-mediated epigenetic plasticity orchestrates tubular regeneration and adaptive repair

Following IR exposure, injured TECs release Damage-Associated Molecular Patterns (DAMPs) into the extracellular space (35,36). These endogenous danger signals activate innate immune pathways - including NF-κB, STAT3, and NLRP3 - thereby triggering pro-inflammatory cascades. Simultaneously, surviving tubular cells initiate repair by undergoing dedifferentiation via a partial EMT program. H2A.J appears essential for coordinating these regenerative responses, as our prior in-vitro and in-vivo data indicate that its radiation-induced incorporation promotes chromatin reprogramming and activates AP1 family transcription factors to drive complex tissue regeneration (23,25,34). In contrast, H2A.J deficiency disrupts the synchronization of this dedifferentiation-redifferentiation cycle. This epigenetic dysregulation, coupled with persistent cellular senescence and chronic inflammation, drives maladaptive repair characterized by progressive tubular atrophy and renal fibrosis. Our current model demonstrates that H2A.J provides the epigenetic plasticity required for the transition from acute injury to functional recovery, whereas its absence accelerates the progression to chronic kidney disease.

### Role of mitochondria in stressed TECs

Proximal TECs are exceptionally vulnerable to genotoxic stress due to their high metabolic demand and near-exclusive reliance on oxidative phosphorylation (6). While mitochondria provide the ATP essential for DNA repair and homeostatic restoration, IR triggers mtDNA damage and the disruption of ion gradients. The subsequent surge in ROS and loss of mitochondrial membrane potential activate apoptotic and necroptotic pathways (37,38). In H2A.J-deficient kidneys, we observed an inherent impairment of mitochondrial function, accompanied by elevated baseline rates of apoptosis and compensatory proliferation. Post-IR, these KO kidneys exhibited a persistent accumulation of damaged, senescent TECs, likely attributable to the failure in replenishing functional TECs due to impaired differentiation. A key driver of this impaired differentiation is likely the bioenergetic collapse - characterized by an inability to restore ATP production, stabilize membranes, or clear damaged organelles via mitophagy - which drives a self-perpetuating cycle of oxidative stress and sterile inflammation. Ultimately, this sustained mitochondrial dysfunction promotes epithelial senescence and tubular atrophy, serving as pivotal mechanism in the transition from acute injury to chronic kidney disease.

### Therapeutic Implications and Future Perspectives

Currently, no targeted therapies exist to address the specific molecular pathways underlying radiotherapy-induced nephrotoxicity (39). A primary challenge remains the identification of interventions capable of reactivating regenerative potential and guiding the coordinated cascade of dedifferentiation, proliferation, and maturation toward functional recovery. Our study identifies histone variant H2A.J as a novel epigenetic regulator of tubular regeneration, offering a potential target to mitigate damage induced by genotoxic stress. Given that the transition from acute injury to chronic kidney disease is driven by sustained inflammation and maladaptive repair, H2A.J represents a high-level regulatory node capable of modulating these multifaceted processes. Consequently, targeting H2A.J-mediated pathways may facilitate the development of innovative therapeutic strategies to preserve renal function during oncological treatments. Since H2A.J is highly expressed in the luminal epithelial cells of various tissues, it may possess similar functions in maintaining barrier integrity across multiple organs.

## MATERIALS AND METHODS

### Mouse Models and Husbandry

To generate H2A.J knockout (KO) mice, a 7-bp deletion was introduced into the *H2afj* locus of the C57BL/6N genome using TALEN-mediated genome editing (Cyagen, Santa Clara, CA, USA). This *H2afj*Δ7 mutation induces a frameshift resulting in a premature stop codon at the beginning of the coding sequence. Homozygous H2A.J KO mice were viable and phenotypically unremarkable, with no significant reproductive or health deficiencies observed under standard conditions. C57BL/6N wild-type (WT) mice (Charles River Laboratories, Sulzfeld, Germany) served as controls and were maintained under identical conditions. All animals were housed in groups in individually ventilated cages (IVC) at 22 ± 1°C and 55 ± 10% humidity, with a 12h light/dark cycle and ad libitum access to standard pellet diet and water. Experimental protocols were approved by the Animal Care and Use Committee of Saarland.

### IR Protocols

WT and KO mice (2–3 months old) were assigned to different IR regimens: Acute whole-body IR: 1x 2 Gy (analyzed 0.5, 5, 24, and 48 hours post-IR) or 1x 10 Gy (analyzed 24 hours post-IR), fractionated whole-body IR: 5x 2 Gy (analyzed 1, 3, and 7 days post-IR) and localized kidney IR: 1x 10 Gy to a small volume (analyzed 1, 3, and 6 months post-IR). Whole-body IR was performed on awake, freely moving animals in Plexiglas containers, whereas mice were anesthetized for localized kidney IR. Plexiglas plates were utilized as build-up material to ensure a homogeneous dose distribution within the renal tissue. 3D dose distributions were calculated using the Eclipse™ treatment planning system (Varian Medical Systems, Palo Alto, CA, USA), confirming dose homogeneity within the renal tissue (Suppl. 1). Irradiations were performed using 6 MeV photons (dose rate: 1 Gy/min) via a TrueBeam™ linear accelerator (Varian Medical Systems). At specified time points, mice were anesthetized (120 µg/g ketamine; 16 µg/g xylazine i.p.) and underwent intracardiac perfusion prior to tissue collection. A minimum of 3-6 animals were analyzed per group and time point.

### Immunofluorescence Microscopy (IFM)

Formalin-fixed, paraffin-embedded (FFPE) kidneys were sectioned (4 µm), deparaffinized in xylene, and rehydrated. Antigen retrieval was performed in boiling citrate buffer, followed by blocking with 10% Immunoblock (Carl Roth, Karlsruhe, Germany). Sections were incubated with primary antibodies against H2A.J (either obtained from ActiveMotif, Waterloo, Belgium, or a custom-made polyclonal antibody raised in guinea pig), 53BP1 (Bio-Techne, Wiesbaden, Germany), Ki67 (Abcam, Cambridge, UK), and Lamin B1 (Abcam). Detection was performed using AlexaFluor-488 or −568 conjugated secondary antibodies (Invitrogen, Waltham, Massachusetts, USA). Slides were mounted in VECTAshield™ containing DAPI (Vector Laboratories, Burlingame, USA).

### Immunohistochemistry (IHC)

Following rehydration and antigen retrieval, sections were incubated with antibodies against H2A.J, phosphorylated H2A.J (custom-made monoclonal antibody raised in rabbit), Caspase-3 (R&D Systems, Nordenstadt, Germany), Megalin (Abcam), TOMM20 (Abcam), KIM-1 (ProteinTech, Planegg-Martinsried, Germany), and Vimentin (Abcam). Biotinylated secondary antibodies (Dako, Glostrup, Denmark) and 3,3′-diaminobenzidine (DAB) were used for visualization. Sections were counterstained with hematoxylin and mounted. Renal iron deposition was assessed using Berlin Blue staining kit (Morphisto, Frankfurt, Germany). To assess renal architecture and quantify collagen deposition as an indicator of fibrosis, longitudinal kidney sections were subjected to Masson’s Goldner Trichrome staining (Morphisto).

### Digital Imaging and Quantitative Analysis

Whole-slide images were acquired at high resolution using a ZEISS Axio Scan.Z1 scanner. Quantitative analysis was performed using HALO® software (Indica Labs, Albuquerque, NM, USA). For each specimen, representative regions of interest (ROIs) were manually annotated, excluding major blood vessels. Analysis modules included: Area Quantification: Berlin Blue and TOMM20. Multiplex IHC: H2A.J and Caspase-3. Random Forest Classifier: H&E based tissue assessment. MiniNet Classifier: Megalin expression. High-plex IF Analysis: 53BP1 and Ki-67. Manual Measurement Tool: Glomerular diameter. Results are expressed as the percentage of positive cells or area relative to the total ROI area/cell count.

### Transmission Electron Microscopy (TEM)

Tissue fragments were fixed in 2.5% glutaraldehyde, post-fixed in 1% osmium tetroxide, and embedded in epoxy resin (EMS, Hatfield, PA, USA). Ultrathin sections (≈70 nm) were contrasted with uranyl acetate and lead citrate and imaged using a Tecnai Biotwin™ TEM (FEI, Eindhoven, Netherlands).

### Scanning Electron Microscopy (SEM)

Cortical blocks (∼1 mm³) were fixed, post-fixed, and dehydrated as described above. Following critical-point drying, samples were sputter-coated with gold and imaged using a JSM-IT210 SEM (Freising, Germany).

### Statistical Analysis

Data were analyzed using GraphPad Prism (v9.4.1) and are presented as mean ± SD (n ≥ 3). Group differences were evaluated via one-way or two-way ANOVA. P-values were defined as: *p < 0.05 (significant), **p < 0.01 (highly significant), and ***p < 0.001 (extremely significant). Significant deviations from non-irradiated controls are indicated by asterisks; differences between experimental groups are indicated by asterisks within brackets.

## Acknowledgments

The authors would like to sincerely thank Prof. D. Fliser (Department of Internal Medicine IV - Nephrology and Hypertension, Saarland University Medical Center, Homburg/Saar, Germany) and Prof. M. van der Laan (Medical Biochemistry & Molecular Biology, Center for Molecular Signalling, Saarland University Medical School, Homburg/Saar, Germany) for their invaluable scientific advice and insightful discussions regarding nephropathology and mitochondrial function.

## Data Availability

The data that support the findings of this study are available from the corresponding author upon reasonable request.

## Supplementary Materials

**Suppl. 1:**
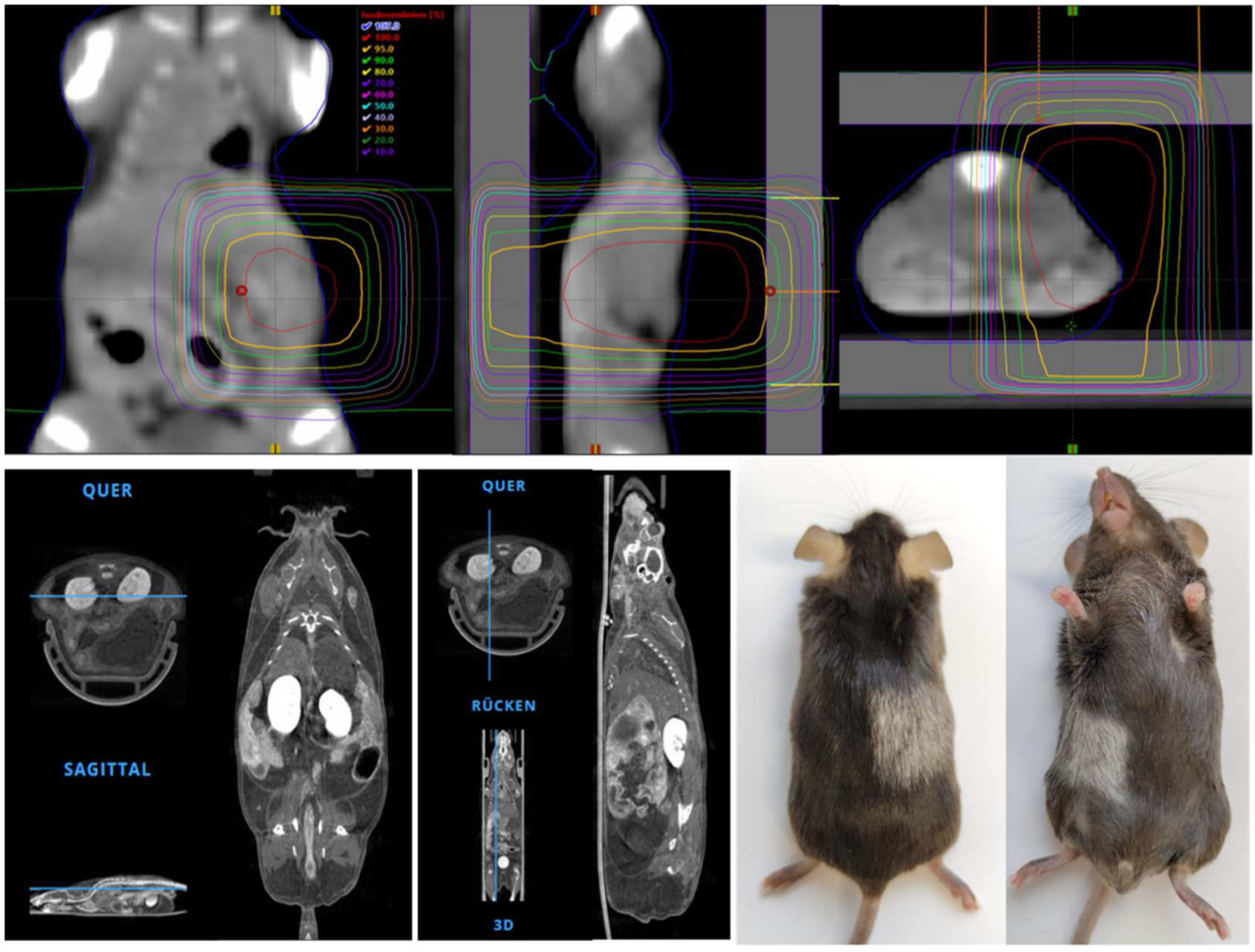
Dose distribution for unilateral kidney irradiation based on CT-guided treatment planning. Graying of the fur within the irradiation field.

**Suppl. 2:**
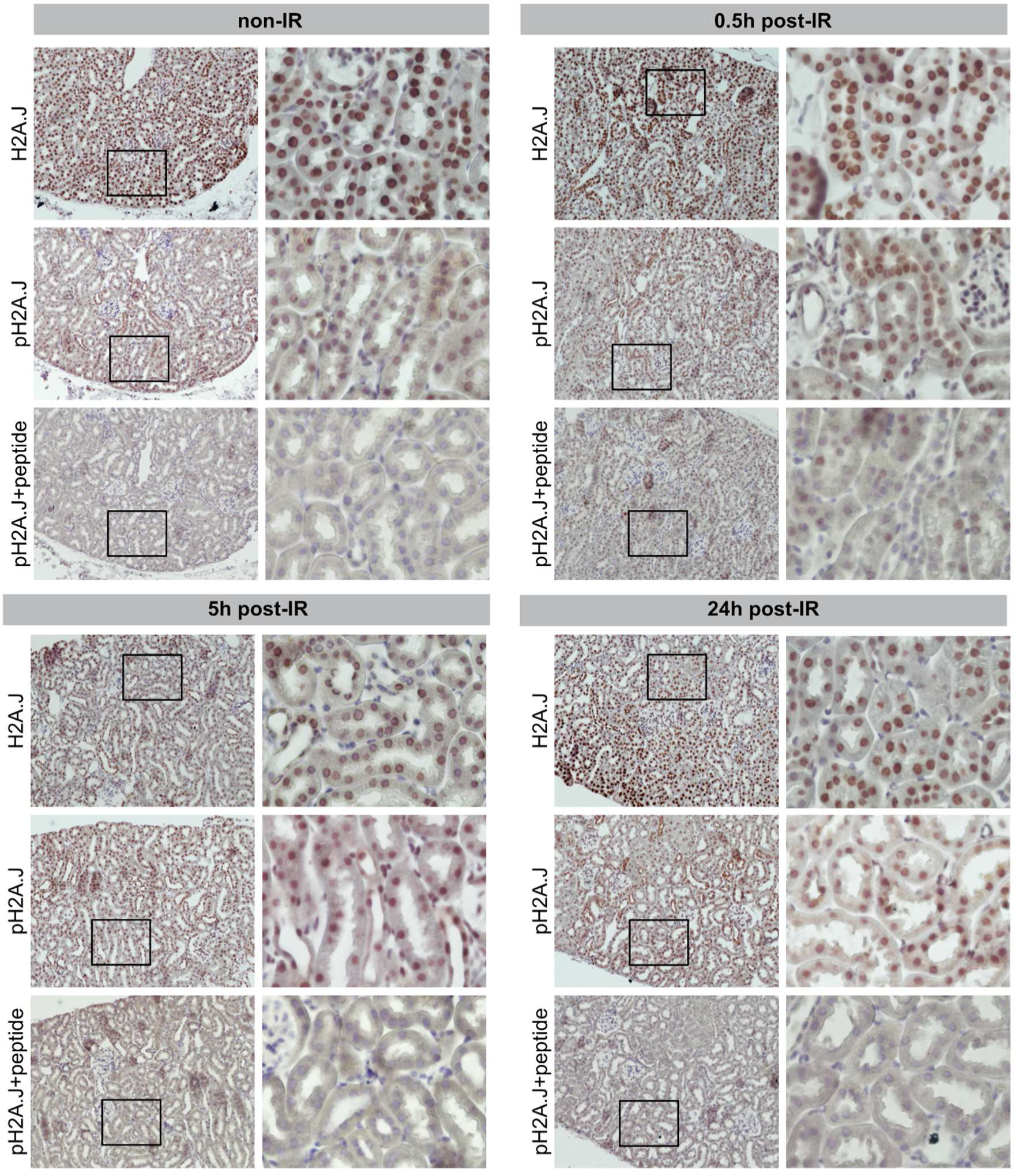
Detection of phosphorylation of H2A.J following IR exposure. Representative ICH micrographs of H2A.J and phosphorylated H2A.J (pH2A.J) staining in WT kidneys at 0.5h, 5h and 24h post-IR compared to non-irradiated controls, with and without peptide treatment.

**Suppl. 3:**
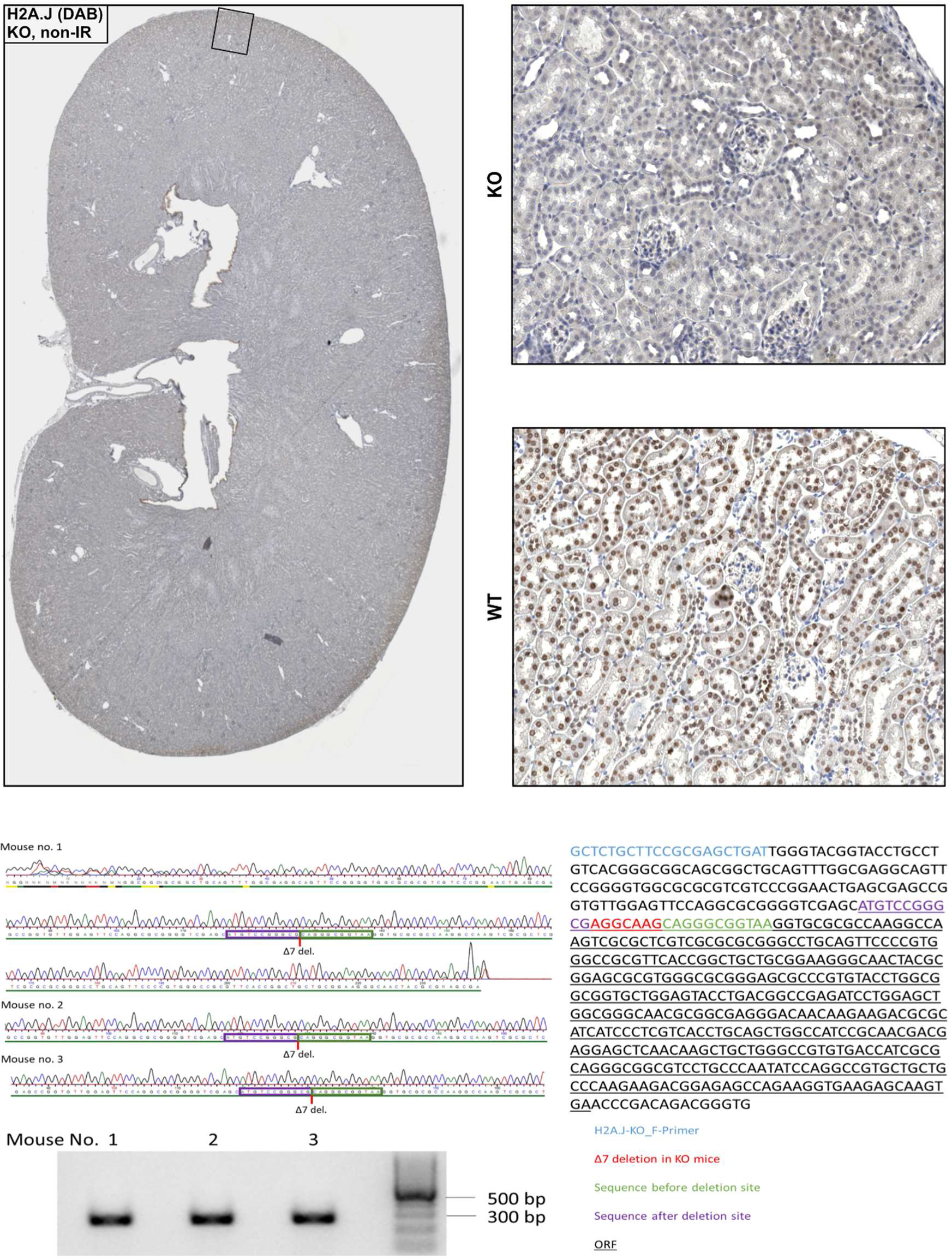
Representative IHC images demonstrating the complete absence of H2A.J expression in the kidneys of KO mice compared to WT littermates. Animal genotypes were verified prior to analysis.

**Suppl. 4:**
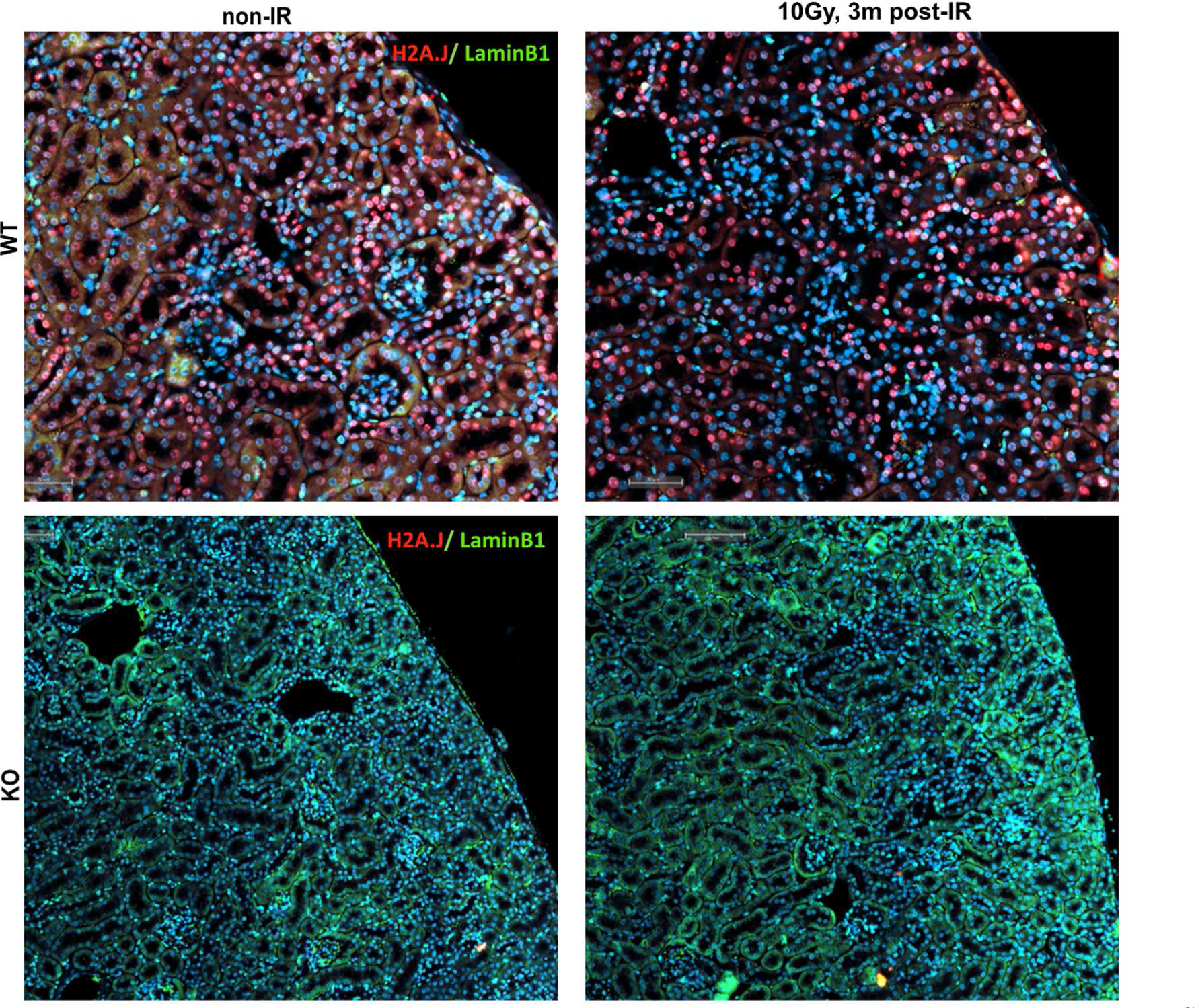
Premature senescence in kidney tissue following IR exposure. Representative IF micrographs of H2A.J-and Lamin B1-stained kidney tissue from WT versus KO mice following IR exposure (10 Gy, 3m post-IR), compared to non-irradiated controls.

**Suppl. 5:**
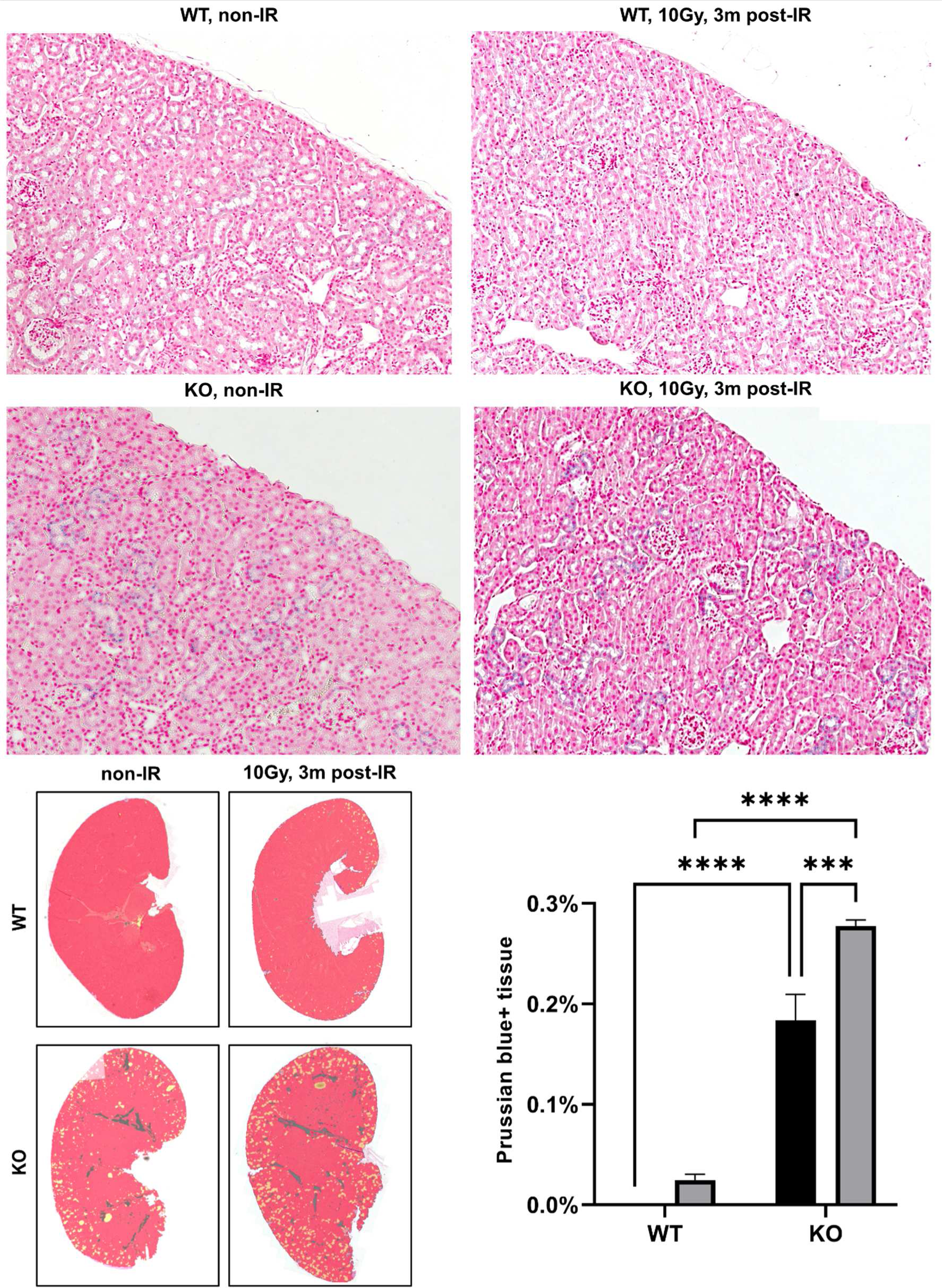
Hemosiderin deposition as an indicator of tubular dysfunction. Representative micrographs of Prussian blue-stained renal tissue from WT and KO mice under baseline conditions and following IR, analyzed by automated image analysis. Data are presented as mean ± SD (n ≥ 3). P-values were defined as: *p < 0.05, **p < 0.01, and ***p < 0.001.

**Suppl. 6:**
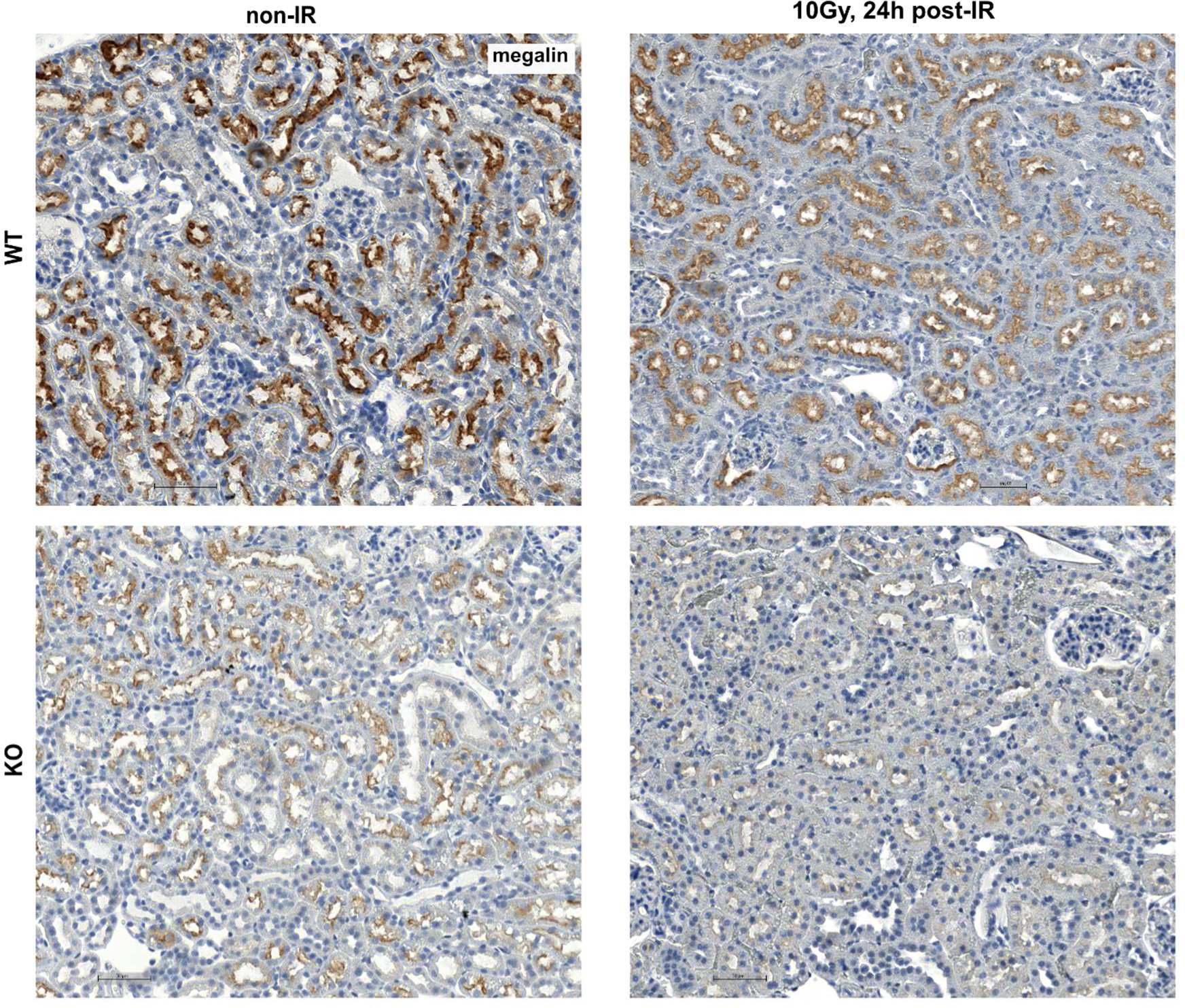
Brush border of TECs an indicator of tubular function. Representative ICH micrographs of megalin-stained WT and KO kidney following IR exposure (10 Gy, 24h post-IR) compared to non-irradiated controls.

**Suppl. 7:**
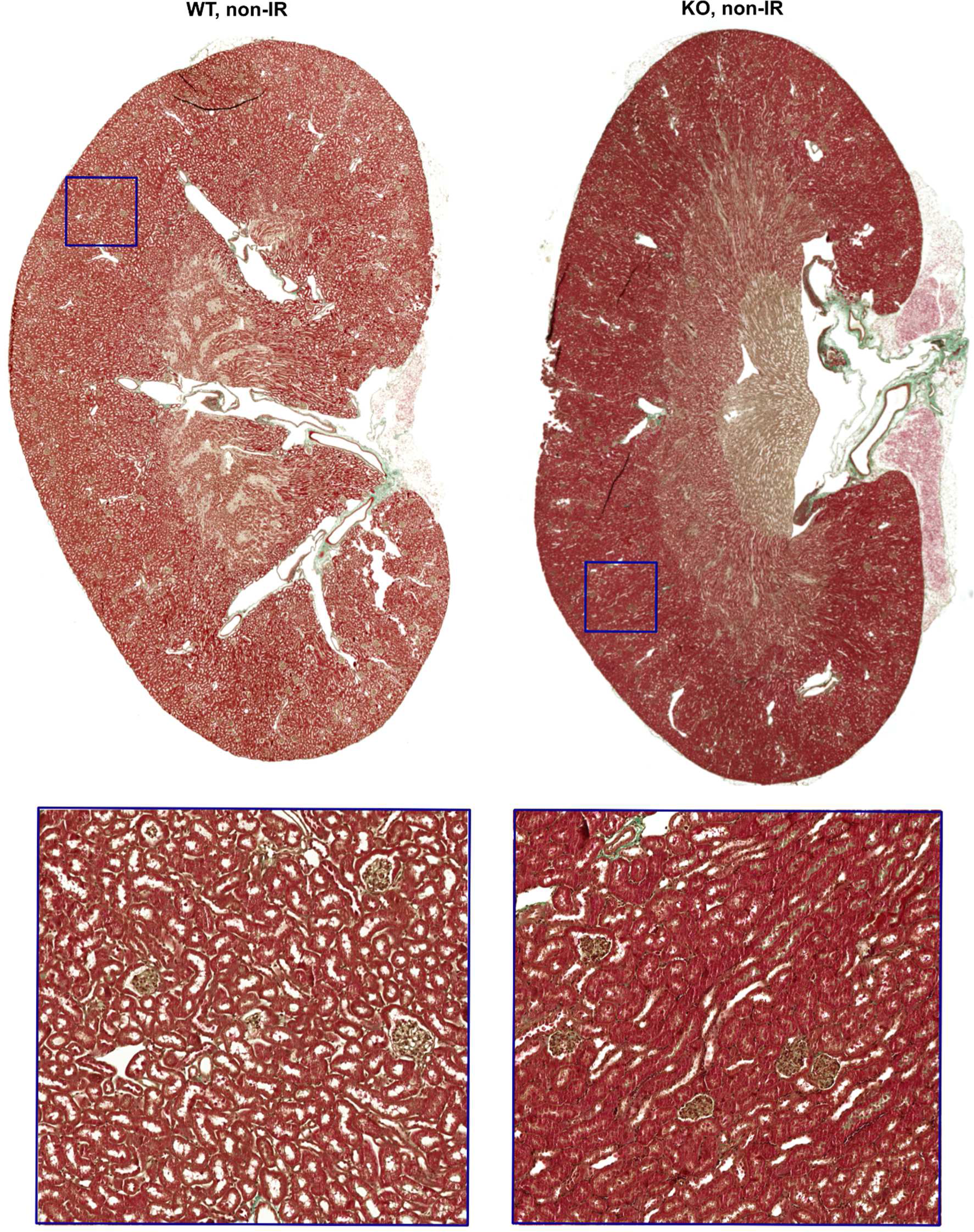
Tubular atrophy in non-irradiated KO kidney. Micrographs of Masson-Goldner-stained non-irradiated WT and KO kidneys. Insets provide higher magnification highlighting tubular atrophy in non-irradiated KO kidney.

**Suppl. 8:**
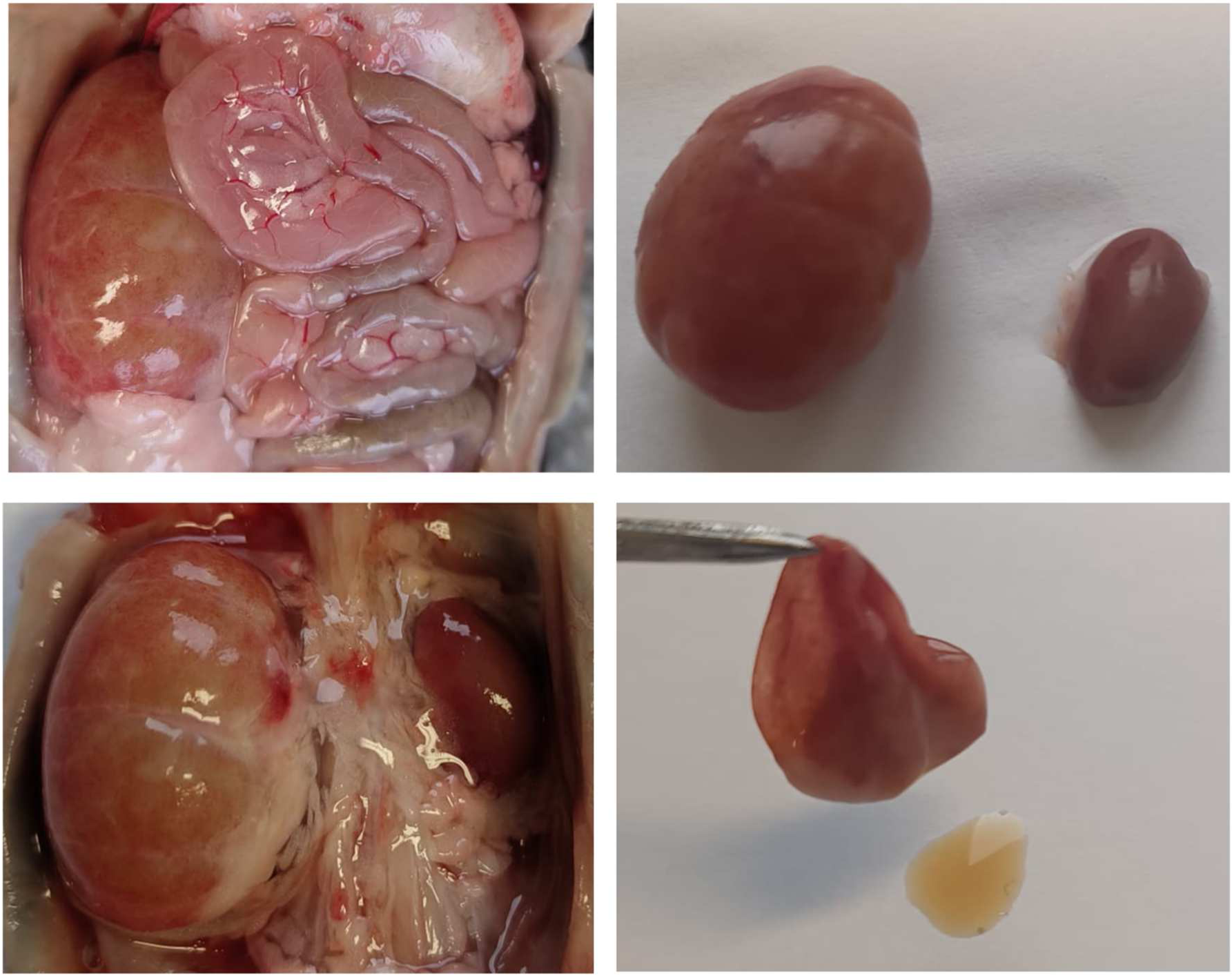
Formation of large cysts in irradiated and aged KO kidneys. Representative photographs of macrocystic kidneys in situ, before and after removal of the intestine (left panels). Excised kidneys before and after aspiration of cyst fluid (right panels).

